# Sarcoidosis resolvers and progressors demonstrate distinct systemic metabolomic profiles

**DOI:** 10.1101/2025.03.10.642370

**Authors:** Bing Ma, Lobelia Samavati, Wonder Puryear Drake

**Affiliations:** University of Maryland School of Medicine, Institute for Genome Sciences, Baltimore, MD, USA; Department of Microbiology and Immunology, University of Maryland School of Medicine, Baltimore, MD, USA; Division of Pulmonary, Critical Care and Sleep Medicine, Department of Medicine, Wayne State University School of Medicine and Detroit Medical Center, Detroit, Michigan; Center for Molecular Medicine and Genetics, Wayne State University School of Medicine, Detroit, Michigan; Division of Infectious Diseases, Department of Medicine, University of Maryland School of Medicine, Baltimore, MD, USA

**Author notes:** **Corresponding Author:** Wonder Drake, MD. **Author Contributions:** Conceptualization: LS, WD; Investigation: LS, BM, WD; Writing of the original draft, review and editing: BM, WD; Visualization: WD; Project administration: WD. **Funding:** National Institute of Health Lung and Blood: HL157533; K24HL127301 (WPD). **Subject Categorization:** 9.39 Sarcoidosis Mechanisms.

**Keywords:** Sarcoidosis, Metabolome, trimethylamine N-oxide

## Abstract

Sarcoidosis is an idiopathic syndrome with striking disparities in clinical outcome. We have previously identified immune distinctions between sarcoidosis clinical cohorts that either resolve or progress. A significantly higher percentage of Programmed Death 1 (PD-1)+Th17 cells was present in sarcoidosis and IPF subjects experiencing disease progression. Metabolites, or bioactive compounds, have been shown to interact with immune cells, such as Th17 cells by regulating cellular processes such as proliferation, signaling and differentiation. Metabolomic analysis was conducted on two subject cohorts: Sarcoidosis (n=19) and healthy controls (n=23). Sarcoidosis subjects were further subdivided by clinical outcome and prior to therapeutic initiation: disease progression or clinical resolution. Metabolites are considered significant if they had a fold change greater than two and a p-value is <0.05 in pairwise 2-sample t-tests. Multiple group comparison was performed using ANOVA and Tukey’s honestly significance difference post hoc tests to identify statistically different pairwise differences. Progressors exhibit significantly elevated serum levels of trimethylamine N-oxide (TMAO) and taurine, with reduced glycerate, alanine, and proline concentrations (Figure 1A-C). TMAO is elevated in sarcoidosis progressors relative to resolvers. Taurine levels are also significantly elevated in progressors, comparing to resolvers. The significant reductions in glycerate, alanine, and proline in progressors compared to resolvers, along with elevated TMAO, point to disruptions in core metabolic pathways, including glycolysis, pyruvate metabolism, and collagen synthesis, that underlie bioenergetic dysfunction, altered amino acid utilization, and impaired tissue repair.

## Introduction

Sarcoidosis is an idiopathic syndrome with striking disparities in clinical outcome. Approximately 60% of patients experience spontaneous resolution of their disease, whereas 40% progress, with 10-20% of sarcoidosis patients ultimately die from lung disease (1). Despite its clinical importance, it remains largely unknown on the mechanisms driving these divergent trajectories. Advances in molecular profiling, such as metabolomics, offers a novel perspective to garner new insights into its cellular mechanisms underlying sarcoidosis pathogenesis.

We have previously identified immune distinctions between sarcoidosis clinical cohorts that either resolve or progress. A significantly higher percentage of Programmed Death 1 (PD-1)+Th17 cells in sarcoidosis and IPF subjects experiencing disease progression (2, 3). Metabolites, or bioactive compounds, have been shown to interact with immune cells, such as Th17 cells by regulating cellular processes such as proliferation, signaling and differentiation (4). Two recent studies employed metabolomic approaches and revealed aberrant metabolic profiles associated with granulomatous inflammation in sarcoidosis compared to healthy controls (5, 6). However, the metabolic distinctions between progressors and resolvers in sarcoidosis remain uncharacterized. Therefore, this study conducted untargeted metabolomic analysis of serum from sarcoidosis patients with disease progression to those who are experiencing resolution.

### Cohort description and data analyses

For study participation, clinical and radiographic criteria were used to define sarcoidosis as has been previously described (5). IRB approval was obtained from Wayne State University and Detroit Medical Center. All investigations with human subjects were conducted according to the principles expressed in the Helsinki Declaration. There were two subject cohorts: Sarcoidosis (n=19) and healthy controls (n=23). Sarcoidosis subjects were further subdivided by clinical outcome: disease progression or clinical resolution. The sarcoidosis patients with progressive disease were characterized by reductions in forced vital capacity (FVC), development of extrapulmonary sarcoidosis, or pulmonary symptom acceleration that necessitates therapeutic intervention. Sarcoidosis subjects who had resolved disease were distinguished by normalized FVC or chest radiograph and resolution of pulmonary symptoms without immunosuppressive therapy. Following written consent, blood samples were obtained from sarcoidosis patients prior to the initiation of any therapeutic agents. ^1^H nuclear magnetic resonance (NMR)-based untargeted metabolomic analysis was used to identify circulating molecules in serum. NMR spectra data processing and calibration is outlined as previously reported (5). Rigorous data pretreatment was performed according to validated procedures (7). Partial least squares discriminant analysis (PLS-DA) was used to demonstrate clusters by groups. Metabolites are considered significant if they show a fold change greater than two and a p-value is <0.05 in pairwise 2-sample t-tests. Multiple group comparison was performed using ANOVA and Tukey’s honestly significance difference post hoc tests to identify statistically different pairwise differences.

## Results and Discussion

Demographically, there were no significant differences by race, sex or age between the sarcoidosis and the healthy control cohorts (p=0.53, two-tailed Fishers). There were also no significant differences by age or sex among sarcoidosis progressors or resolvers; there was a higher percentage of Caucasians in the resolver cohort compared to progressors (p=0.015, two tailed Fishers). Metabolomic analysis demonstrates distinctions between healthy controls and sarcoidosis patients (Table 1), with further differentiation associated with sarcoidosis disease severity (Table 1). Progressors exhibit significantly elevated levels of trimethylamine N-oxide (TMAO) and taurine, with reduced glycerate, alanine, and proline concentrations (Figure 1A-C). TMAO is elevated in sarcoidosis progressors relative to resolvers. It has a well-established pro-inflammatory profile relevant to granulomatous inflammation, including NLRP3 inflammasome activation, endothelial dysfunction induction, and oxidative stress augmentation (8). Notably, TMAO enhances Th17-mediated inflammatory responses, as demonstrated by increased Th17 cells recruitment in autoimmune murine models (9), suggesting a metabolic basis for the observed immunological dysregulation. TMAO is synthesized in the liver via the oxidation of circulating trimethylamine (TMA), a metabolite produced exclusively by the gut microbiota metabolizing dietary substrates such as L-carnitine and choline that are abundant in red meat, fish, and dairy products (8). The parallel changes in carnitine and choline levels in sarcoidosis patients compared to healthy controls further supports a metabolic axis that links dietary substrate availability, gut microbial metabolism, and systemic inflammation. Further investigation of TMA-producing gut microbiota in sarcoidosis may clarify how dietary patterns influence disease progression and reveal novel targets for monitoring and therapeutic intervention.

**Table 1.** Multiple Comparison on serum metabolites from sarcoidosis progresser, sarcoidosis resolver and healthy control cohorts.

| Metabolite | Multiple group comparison* |  | Progressor and resolver comparison |  |
| --- | --- | --- | --- | --- |
|  | FDR | significant comparison | t.stat | p value |
| Glycerate | 1.30E-06 | control - resolver; control - progressor; resolver - progressor | 3.719 | 0.002 |
| Homocysteine | 1.57E-06 | resolver - control; progressor - control |  |  |
| Cystine | 1.64E-05 | resolver - control; progressor - control |  |  |
| Aminobutyrate | 0.001 | control - resolver; control - progressor |  |  |
| TMAO | 0.006 | progressor - control | -2.1107 | 0.049 |
| Alloisoleucine | 0.006 | control - resolver; control - progressor |  |  |
| Glycine | 0.011 | resolver - control; progressor - control |  |  |
| Carnitine | 0.012 | progressor - control |  |  |
| Taurine | 0.012 | progressor - control; progressor - resolver | -2.9883 | 0.008 |
| Succinate | 0.014 | control - resolver; control - progressor |  |  |
| Glutamine | 0.014 | control - resolver; control - progressor |  |  |
| Isoleucine | 0.016 | control - resolver; control - progressor |  |  |
| Pyruvate | 0.016 | resolver - control; progressor - control |  |  |
| Choline | 0.017 | control - progressor |  |  |
| Hydroxybutyrate | 0.017 | resolver - control |  |  |
| Alanine | 0.045 | control - progressor | 2.2361 | 0.038 |
| Propylene.glycol | 0.045 | control - progressor |  |  |
| Proline |  |  | 2.2255 | 0.039 |

\* multiple group comparison was done using Fisher's LSD, 2-sample t-tests used in pairwise comparison

Taurine levels are also significantly elevated in progressors, comparing to resolvers (Table 1). Given that taurine is produced downstream of cysteine in the transsulfuration pathway (5), its increase indicates dysregulation of sulfur amino acid metabolism and enhanced oxidative stress. Prior studies have demonstrated reduced glutathione levels in sarcoidosis patients, suggesting impaired antioxidant capacity (10). Consequently, this metabolic profile implies a shift wherein cysteine is preferentially channeled toward taurine synthesis rather than glutathione production, potentially serving as an adaptive response to oxidative damage and modulate immune activation. Moreover, the significant reductions in glycerate, alanine, and proline in progressors compared to resolvers, along with elevated TMAO (Figure 1), point to disruptions in core metabolic pathways, including glycolysis, pyruvate metabolism, and collagen synthesis, that underlie bioenergetic dysfunction, altered amino acid utilization, and impaired tissue repair.

**Figure 1.**
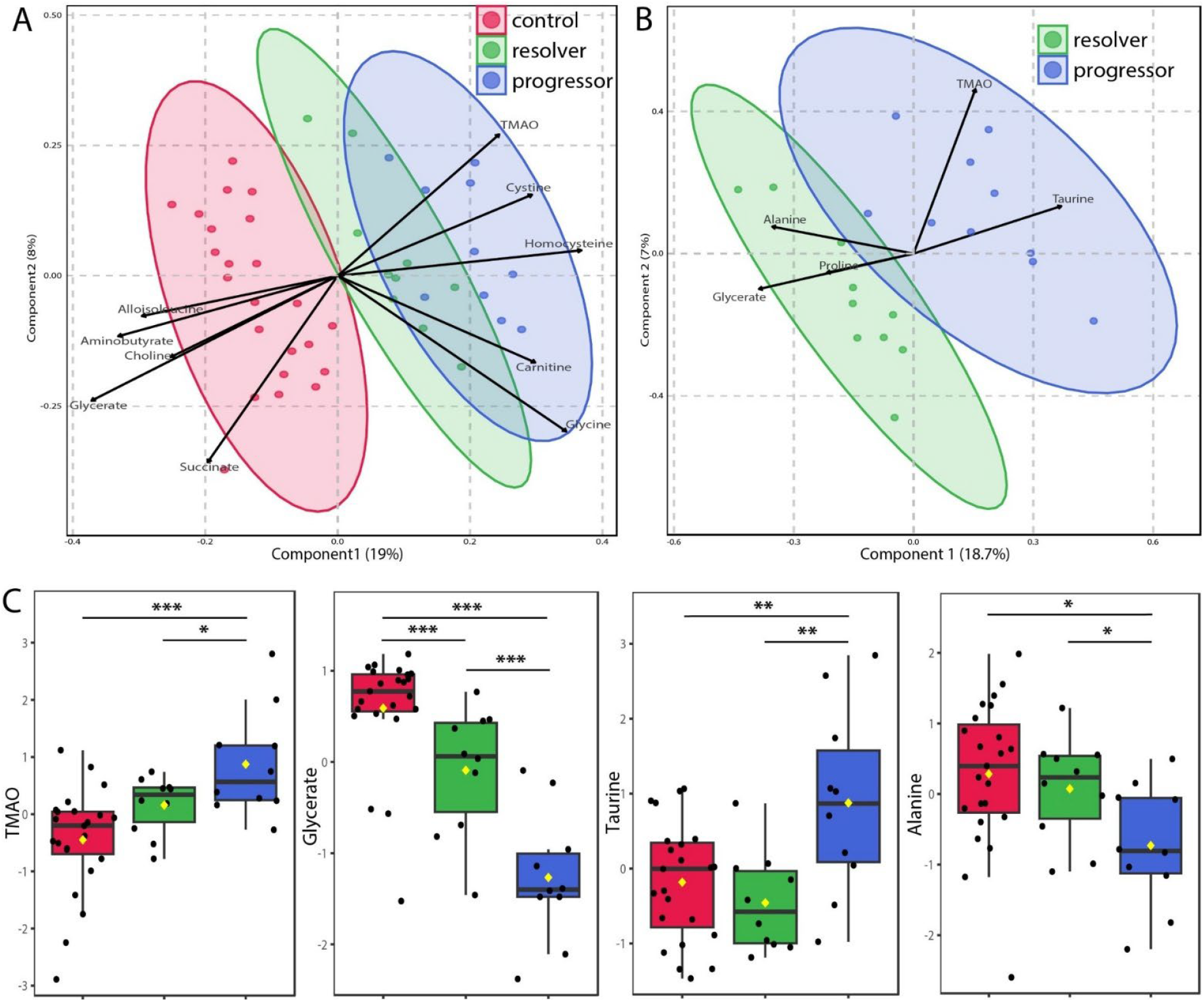
Comparison of metabolome clusters among sarcoidosis and healthy control cohorts. **A**) PLS-DA demonstrates clusters among sarcoidosis resolver, progressor, and controls, and **B**) only between resolver and progressor. Circles indicate 95% confidence region of each group. Distinct signatures exemplified using boxplots **C**). Metabolites with a fold change greater than two and a p-value is <0.05 in pairwise 2-sample t-tests are considered significant. Multiple group comparison were performed using ANOVA and Tukey’s honestly significance difference post hoc tests to identify statistically different pairwise differences. *p<0.05, **p<0.01, ***p<0.001.

Despite the insights from this metabolomic investigation, several limitations warrant consideration. This pilot study was conducted with a relatively small cohort. Validation in larger, more diverse cohorts using advanced metabolomic technologies with more comprehensive coverage is needed to confirm the observed signatures. The cross-sectional nature of the study limits causal inference; longitudinal investigations tracking metabolite changes throughout disease progression or resolution would provide valuable insights. Additionally, while the data suggest connections between metabolic alterations and immune dysregulation, a more integrative approach incorporating parallel immunophenotyping, microbiome characterization, and metabolomics would better elucidate the complex interplay between microbial dysbiosis, metabolite production, and sarcoidosis immune regulation. Such multidimensional analyses will ultimately reveal novel therapeutic targets addressing the multifaceted etiology of sarcoidosis progression.

## References

1. Crouser ED, Maier LA, Wilson KC, Bonham CA, Morgenthau AS, Patterson KC, Abston E, Bernstein RC, Blankstein R, Chen ES, Culver DA, Drake W, Drent M, Gerke AK, Ghobrial M, Govender P, Hamzeh N, James WE, Judson MA, Kellermeyer L, Knight S, Koth LL, Poletti V, Raman SV, Tukey MH, Westney GE, Baughman RP. Diagnosis and Detection of Sarcoidosis. An Official American Thoracic Society Clinical Practice Guideline. Am J Respir Crit Care Med 2020; 201: e26–e51.

2. Celada LJ, Kropski JA, Herazo-Maya JD, Luo W, Creecy A, Abad AT, Chioma OS, Lee G, Hassell NE, Shaginurova GI, Wang Y, Johnson JE, Kerrigan A, Mason WR, Baughman RP, Ayers GD, Bernard GR, Culver DA, Montgomery CG, Maher TM, Molyneaux PL, Noth I, Mutsaers SE, Prele CM, Peebles RS, Jr., Newcomb DC, Kaminski N, Blackwell TS, Van Kaer L, Drake WP. PD-1 up-regulation on CD4(+) T cells promotes pulmonary fibrosis through STAT3-mediated IL-17A and TGF-beta1 production. Sci Transl Med 2018; 10.

3. Chioma OS, Mallott EK, Chapman A, Van Amburg JC, Wu H, Shah-Gandhi B, Dey N, Kirkland ME, Blanca Piazuelo M, Johnson J, Bernard GR, Bodduluri SR, Davison S, Haribabu B, Bordenstein SR, Drake WP. Gut microbiota modulates lung fibrosis severity following acute lung injury in mice. Commun Biol 2022; 5: 1401.

4. Tannahill GM, Curtis AM, Adamik J, Palsson-McDermott EM, McGettrick AF, Goel G, Frezza C, Bernard NJ, Kelly B, Foley NH, Zheng L, Gardet A, Tong Z, Jany SS, Corr SC, Haneklaus M, Caffrey BE, Pierce K, Walmsley S, Beasley FC, Cummins E, Nizet V, Whyte M, Taylor CT, Lin H, Masters SL, Gottlieb E, Kelly VP, Clish C, Auron PE, Xavier RJ, O’Neill LA. Succinate is an inflammatory signal that induces IL-1beta through HIF-1alpha. Nature 2013; 496: 238–242.

5. Geamanu A, Gupta SV, Bauerfeld C, Samavati L. Metabolomics connects aberrant bioenergetic, transmethylation, and gut microbiota in sarcoidosis. Metabolomics 2016; 12.

6. Rai MK, Yadav S, Jain A, Singh K, Kumar A, Raj R, Dubey D, Singh H, Guleria A, Chaturvedi S, Khan AR, Nath A, Misra DP, Agarwal V, Kumar D. Clinical metabolomics by NMR revealed serum metabolic signatures for differentiating sarcoidosis from tuberculosis. Metabolomics 2023; 19: 92.

7. van den Berg RA, Hoefsloot HC, Westerhuis JA, Smilde AK, van der Werf MJ. Centering, scaling, and transformations: improving the biological information content of metabolomics data. BMC Genomics 2006; 7: 142.

8. Koeth RA, Wang Z, Levison BS, Buffa JA, Org E, Sheehy BT, Britt EB, Fu X, Wu Y, Li L, Smith JD, DiDonato JA, Chen J, Li H, Wu GD, Lewis JD, Warrier M, Brown JM, Krauss RM, Tang WH, Bushman FD, Lusis AJ, Hazen SL. Intestinal microbiota metabolism of L-carnitine, a nutrient in red meat, promotes atherosclerosis. Nat Med 2013; 19: 576–585.

9. Gonzalez-Correa C, Moleon J, Minano S, Visitacion N, Robles-Vera I, Gomez-Guzman M, Jimenez R, Romero M, Duarte J. Trimethylamine N-Oxide Promotes Autoimmunity and a Loss of Vascular Function in Toll-like Receptor 7-Driven Lupus Mice. Antioxidants (Basel) 2021; 11.

10. Boots AW, Drent M, Swennen EL, Moonen HJ, Bast A, Haenen GR. Antioxidant status associated with inflammation in sarcoidosis: a potential role for antioxidants. Respir Med 2009; 103: 364–372.

